# DAESC+: High-performance, integrated software for single-cell allele-specific expression data

**DOI:** 10.1101/2025.09.03.674100

**Authors:** Tengfei Cui, Guanghao Qi

## Abstract

Single-cell allele-specific expression (ASE) provides valuable insights into gene regulatory mechanisms. However, its utility is limited by the lack of dedicated computational tools. We present DAESC+, an end-to-end software package for the processing and analysis of single-cell ASE. The preprocessing module, DAESC-P, is a user-friendly bioinformatics pipeline to obtain ASE counts from multiplexed scRNA-seq data. The analysis module, DAESC-GPU, is a scalable tool for differential ASE analysis powered by graphics processing units (GPUs). We demonstrated that DAESC-P is more accurate than the existing SALSA pipeline. DAESC-GPU is dozens of times faster than our previous method (DAESC) and scalable to over a million cells. Applying DAESC+ to a subset of the OneK1K cohort, we identified 15 genes exhibiting differential regulatory patterns between naïve and central memory CD4+ T cells, and 2 genes between naïve and memory B cells.

## Introduction

Allele-specific expression (ASE) is a powerful signal to study multiple regulatory mechanisms^1–4^. ASE is typically quantified from RNA-sequencing (RNA-seq) by the number of reads from two parental alleles at an exonic single-nucleotide polymorphism (SNP). In the past decade, single-cell RNA-seq (scRNA-seq) has advanced the quantification of ASE to single-cell resolution. Single-cell ASE was used to study a range of biological effects, such as transcriptional bursting^5,6^, genomic imprinting^7^, cis-regulatory effects^8–11^, and cancer clonal evolution^12^. With the growth of single-cell data and emerging technologies such as long-read sequencing, ASE has the potential to lead to many novel discoveries^4,13,14^.

Despite the importance of single-cell ASE, only a few statistical methods are available for the analysis of such data^5,7,15,16,9–11,4^. For example, SCALE^5^ characterizes random monoallelic expression and estimates transcriptional bursting kinetics. scDALI^9^ tests for homogeneous and heterogeneous allelic imbalance across cell states. Airpart^10^ aims to cluster genes and cells based on patterns of allelic imbalance. We previously developed DAESC^11^, a method for differential ASE (D-ASE) analysis using scRNA-seq data of multiple individuals. DAESC can be applied to any comparison of interest, such as discrete cell types, continuous cell states, case-control disease status, etc. Through simulation studies and read data applications, DAESC was demonstrated to have robust type I error and high power, and identified hundreds of genes that are dynamically regulated during endoderm differentiation^11^. However, the variational EM algorithm adopted by DAESC is computationally intensive. The original R implementation does not scale well to large datasets containing more than a million cells. With rapid expansion of single-cell RNA-seq data to population level^17^, DAESC has the potential for many new applications, but its utility is limited by lack of computational efficiency.

In addition, there is no consensus bioinformatics pipeline to obtain single-cell ASE counts from raw scRNA-seq reads. This task involves several complex steps of bioinformatics processing, including read mapping^18^, genotype calling^19^, haplotype phasing^20^, mapping bias removal^21^, and ASE read counting^19^. Each step requires a specialized software which needs to be configured individually. Most studies developed a custom pipeline specifically for the dataset being analyzed, which may not be applicable to other datasets^22,5,6,8^. SCReadCounts^23^, a generalizable tool, handles only ASE read counting but no other steps. It takes as input single-nucleotide variants (SNVs) and BAM files, which often require significant effort and expertise to generate. Software has also been developed to handle the HLA region specifically^24^. A recent package, SALSA^25^, provides an end-to-end pipeline to obtain allele-specific counts from 10x Genomics data. However, SALSA does not account for multiplexing (multiple samples are sequenced together) and assumes that all reads from a FASTQ file are from the same individual. This can lead to erroneous genotype calls and ASE counts due to mixing reads from different individuals. In addition, SALSA requires a complex computing environment comprised of dependency packages and large reference datasets, which can be difficult to configure. Users would have to peruse the documentation of other software to identify the correct components.

We present DAESC+, an integrated software package for the processing and analysis of single-cell ASE data. The preprocessing module of DAESC+ is a user-friendly bioinformatics pipeline to obtain ASE counts from raw reads of 10x Genomics scRNA-seq data (DAESC-P). DAESC-P allows multiplexing, requires minimal user input, and reduces workload of environment configuration. The analysis module is a high-performance tool for single-cell differential ASE analysis implemented using graphics processing units (DAESC-GPU). DAESC-GPU was shown to be dozens of times faster than DAESC and can be applied to over 1 million cells. Application of DAESC+ to the scRNA-seq data of the OneK1K cohort identified genes that have distinct patterns of allelic imbalance between naïve and memory T cells and B cells.

## Results

### Overview of DAESC+

DAESC+ is comprised of DAESC-P for preprocessing and DAESC-GPU for differential analysis. DAESC-P begins with an integrated script to download sequencing data from the Sequence Read Archive (SRA)^26,27^ with sample IDs as input, convert SRA files to FASTQ files and rename them with 10x naming convention. Next, if scRNA-seq data are multiplexed, i.e., multiple samples are sequenced together, DAESC-P splits each FASTQ file by individual and combines individual-specific FASTQ files from all sequencing runs into a single file (**Figure 1**). All FASTQ files will then be aligned to the genome using Cell Ranger (version 9.0.1, 10x Genomics), with path of the files and list of individuals as input. We then adopted the SALSA pipeline^25^ for the following steps, including genotype calling and quality control, haplotype phasing, alignment bias removal by WASP^21^, and ASE read counting. Although DAESC-P builds upon SALSA, its additional functions to download data and handle multiplexing makes it applicable to a wider range of datasets. It integrates many commands and allows users to configure environment and run complex analysis with a few lines of code. In contrast, SALSA requires users to configure many dependencies separately.

**Figure 1.**
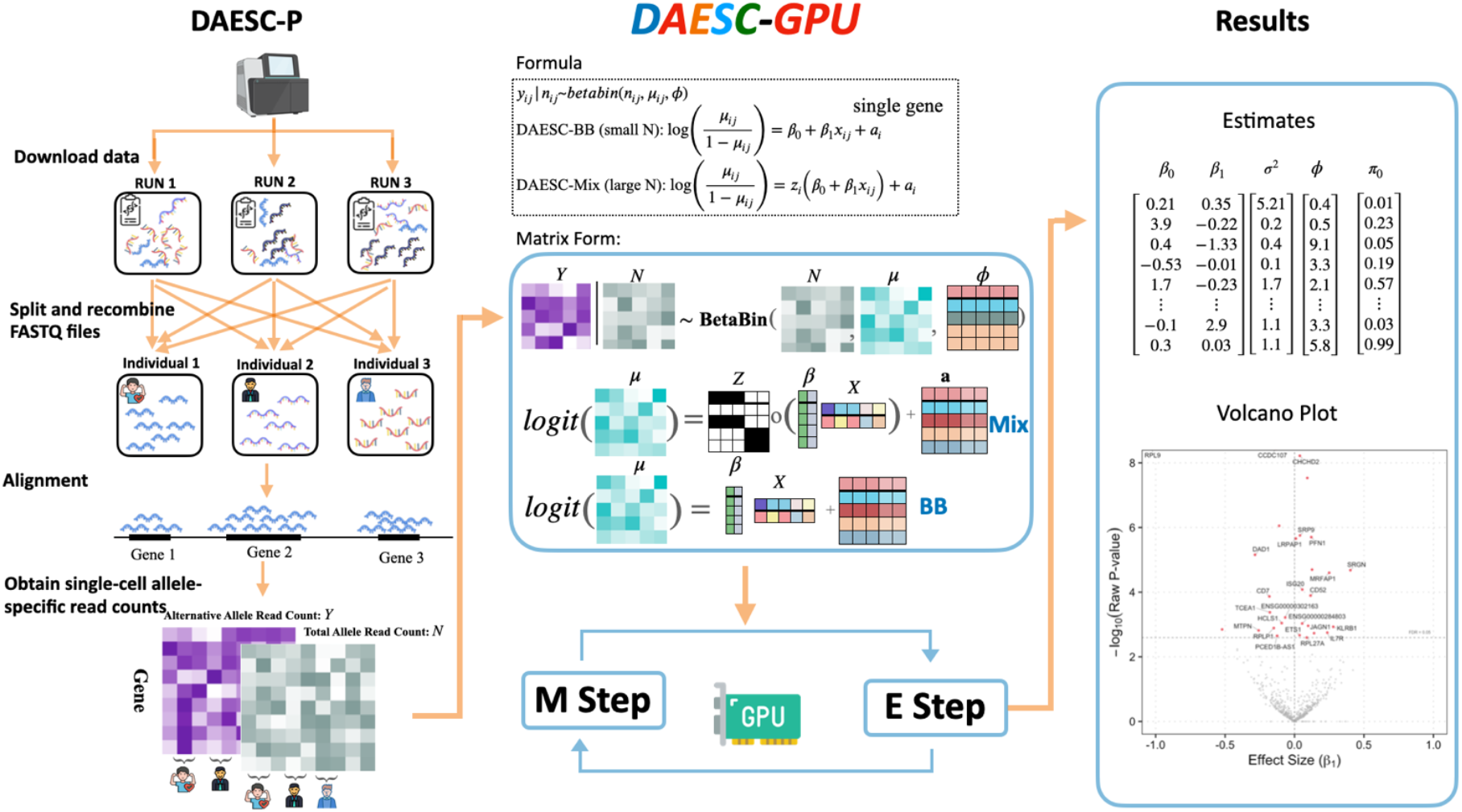
Schematic of DAESC+. DAESC+ is an end-to-end package for processing and analysis of single-cell ASE data, comprised of two modules. 1) DAESC-P is a preprocessing module to obtain single-cell ASE counts from raw reads of multiplexed scRNA-seq data. 2) DAESC-GPU is a scalable tool for single-cell differential allele-specific expression (D-ASE) analysis, leveraging large matrix computation and massive parallel computing using graphics processing units (GPUs).

DAESC-GPU is a scalable re-implementation of DAESC using GPUs. The original DAESC adopts the same beta-binomial regression model with allele-specific counts as outcome and any condition of interest as predictor (see **Methods** for summary)^11^. It incorporates individual-specific random effects to account for sample repeat structure arising from having multiple cells per individual, and latent variables to conduct implicit haplotype phasing. In contrast to the DAESC approach of analyzing one gene at a time, DAESC-GPU analyzes all genes simultaneously using large matrix operations (**Figure 1**). These operations are implemented in CuPy, a Python library using CUDA Toolkit, which dramatically accelerates computation using GPUs through massive parallel processing. Since there is no efficient built-in optimizer, we wrote custom functions to implement a fully vectorized BFGS algorithm across all genes. We used kernels in CuPy, which run directly on GPUs, to integrate multiple mathematical operations into one single specialized function, achieving much higher efficiency than built-in CuPy functions. Our software can be run on GPUs of Google Colab free of charge or other GPUs for higher performance. See **Methods** for details.

### Performance of DAESC-GPU on endoderm differentiation data

To evaluate the performance of DAESC-GPU, we conducted differential ASE analysis along pseudotime using single-cell RNA-seq data (Smart-seq2) from an endoderm differentiation experiment^8^ (see **Methods** for details). ASE count data were provided by the original study and hence DAESC-P did not need to be applied. After processing as described in our earlier paper^11^, we obtained allele-specific read counts for 4,102 genes and 30,474 cells collected from 105 individuals. DAESC-GPU was 30∼90 times faster than DAESC for analyzing this dataset (**Figure 2a**). DAESC-GPU-BB used 64 minutes on a T4 GPU and 32 minutes on an A100 GPU, while DAESC-BB used 1,755 minutes on CPUs from Google Colab and 1,866 minutes on the UW Biostatistics Computing Cluster (referred to as “UW Server”). The patterns for the DAESC-GPU-Mix and DAESC-Mix were similar, though both methods used longer time than the BB version due to the extra computational complexity (**Figure 2a**). DAESC-GPU produced nearly identical estimates to DAESC. The Kendall correlation between DAESC-GPU and DAESC estimates were higher than 0.9 for *β*_0_, *β*_1_, σ^2^, and ϕ for both the BB and Mix model (**Figure 2b, Supplementary Figures 1 and 2**). The correlation was weaker for π_0_ for Mix (*r* = 0.69, Supplementary **Figure 2**). However, this parameter is of secondary importance and does not directly impact the interpretation of the model. For hypothesis testing of differential ASE, DAESC-GPU and DAESC detected nearly identical sets of dynamic ASE genes at varying FDR thresholds (**Figure 2c**), especially for the BB model. There is slight discrepancy in the sets detected by DAESC-Mix and DAESC-GPU-Mix, but the overlap is still above 90%.

**Figure 2.**
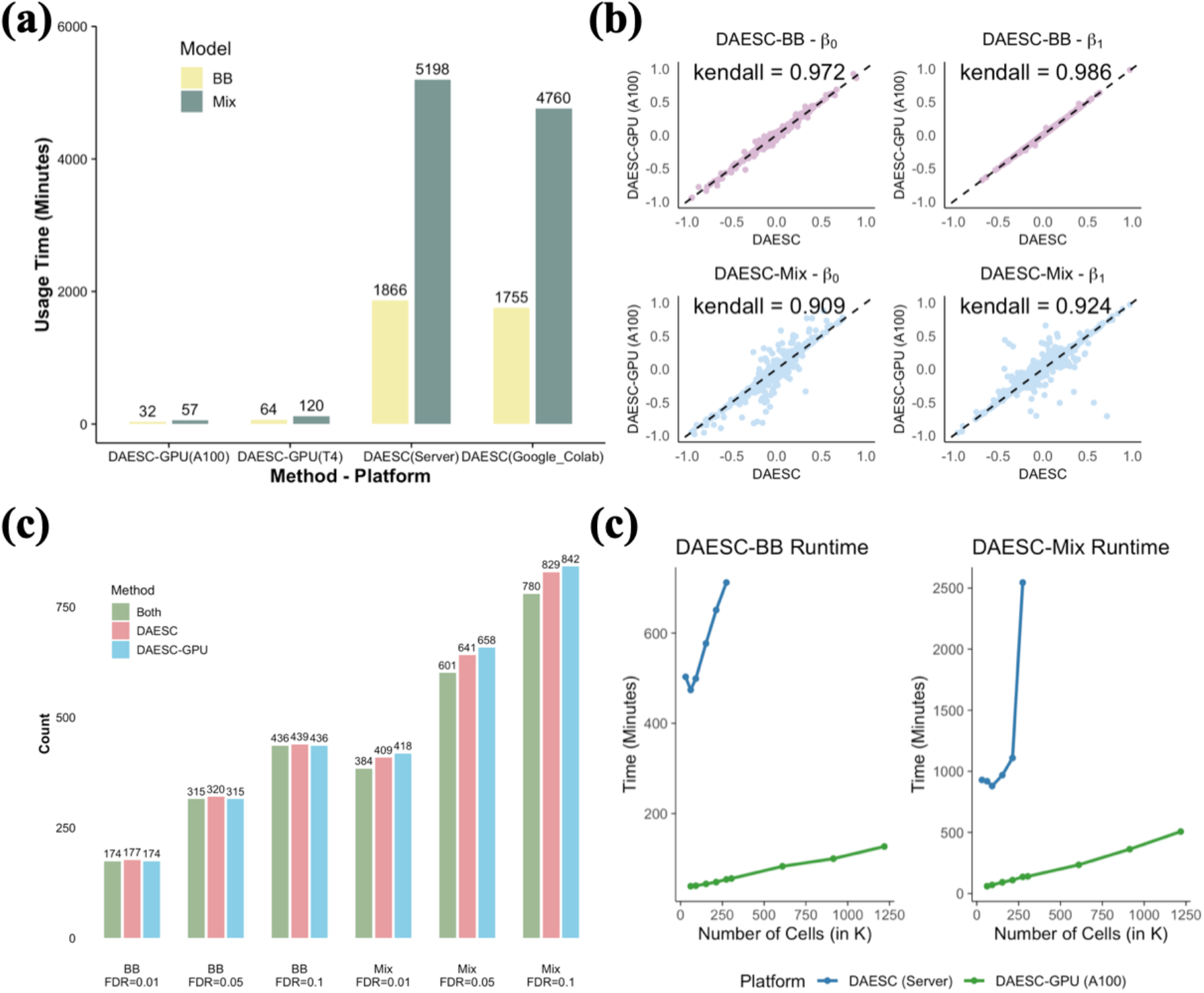
The performance of DAESC-GPU on endoderm differentiation data. (a) Runtime of DAESC-GPU (A100 and T4) vs DAESC R package (UW Server and Google Colab, 8 CPUs) for D-ASE along pseudotime for endoderm differentiation data. (b) Comparison of the estimates of main model parameters between DAESC-GPU and DAESC. Each dot is a gene. Black dashed line is *y* = *x*. Correlation (Kendall’s τ) of the estimate between two methods is marked in the figure. (c) Number of significant genes identified by DAESC, DAESC-GPU and both under varying FDR thresholds. (d) Runtime of DAESC-GPU vs DAESC (R package, 8 CPUs) with varying number of cells in simulation studies.

Next, we tested the scalability of DAESC-GPU in a simulation study (see **Methods** for details). The runtime of DAESC increased rapidly as the number of cells increased, while that of DAESC-GPU increased only moderately and did not present major challenges for analyzing >1 million cells and 2,000 genes (**Figure 2d**). For example, DAESC-BB required 757 minutes to analyze 0.27 million cells, whereas DAESC-GPU-BB completed the analysis of 1.2 million cells in only 127 minutes. Similarly, DAESC-Mix used 2,545 minutes (1.77 days) to analyze 0.27 million cells, while DAESC-GPU-Mix needed only 457 minutes to analyze 1.2 million cells. These results demonstrated that DAESC-GPU is many times faster than DAESC and scalable to large single-cell ASE datasets.

### Processing and analysis of the OneK1K data

We applied DAESC+ to the OneK1K cohort^17^ which was comprised of 982 donors of Northern European ancestry. Multiplexed scRNA-seq data were collected using 10x Genomics Chromium for ∼1.27 million peripheral blood mononuclear cells (PBMCs). Due to large computational demand for processing raw sequencing data, we restricted our analysis to pools 1-10 (out of 77) comprised of 104 individuals. We used DAESC-P to download and process raw reads from SRA and obtained ASE read counts. For benchmarking, we also applied SALSA^25^ to conduct the same processing. SALSA detected non-zero ASE counts in all variant-individual pairs among the top 30 highly expressed variants, even though some of the variants were homozygous in some individuals (**Figure 3**). This is because SALSA conducted genotype calling using multiplexed FASTQ files, mixing reads from multiple individuals and falsely calling homozygotes as heterozygotes. This error led to false monoallelic expression for many individuals, severely biasing the results (**Supplementary Figure 3**). In contrast, DAESC-P correctly detected zero ASE counts where the genotypes are homozygous by splitting and recombining multiplexed FASTQ files based on individuals (**Figure 3**).

**Figure 3.**
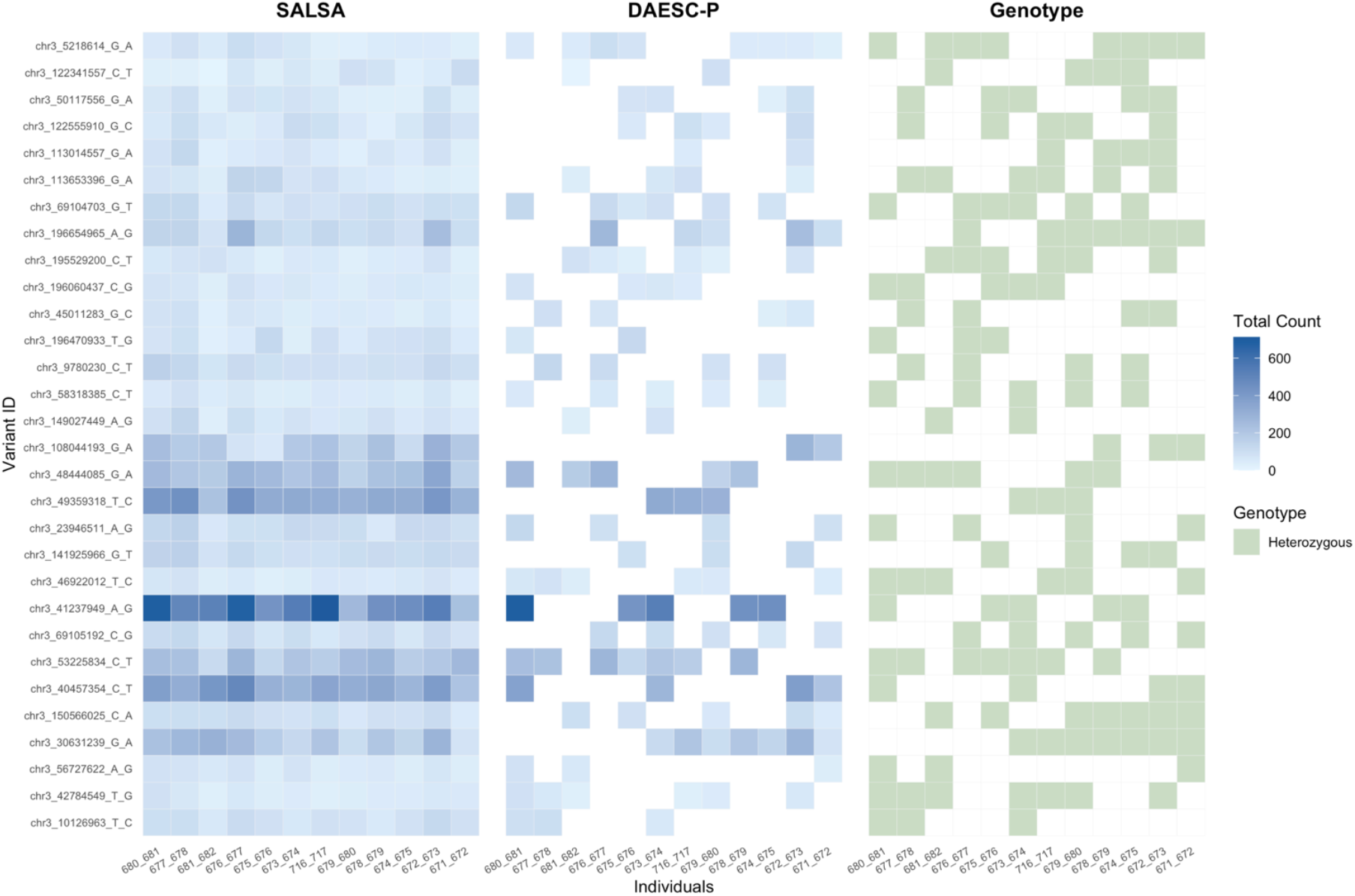
Comparison of ASE counts quantified by DAESC-P and SALSA. The Y-axis shows the top 30 variants of highest total allele-specific read depth quantified by DAESC-P, and the X-axis shows the 12 individuals in Pool 4. Left: total allele-specific read counts quantified by SALSA. Middle: total allele-specific read counts quantified by DAESC-P. White squares represent zero counts (left and middle). Right: heterozygosity of the variants for each individual. Heterozygous genotypes are colored green. Homozygous or uncalled genotypes are colored white.

After preprocessing, we obtained single-cell ASE data of 160,709 cells (See **Methods** for details) from OneK1K. The dataset contained 29 cell types (**Figure 4a**). We used DAESC-GPU to test differential ASE for three immune cell types between naïve versus memory subtypes: CD4+ T cells (naïve and central memory), CD8+ T cells (naïve and central memory), and B cells (naïve and memory). They were among some of the most abundant cell types in the sample (**Figure 4b**). Despite the sparsity of the data, DAESC-GPU remained over 10 times faster than DAESC for both the BB and Mix models in most scenarios (**Figure 4c**). For example, DAESC-GPU-Mix completed D-ASE analysis between naïve versus memory CD4+ naïve T cells in 7.9 minutes (A100) and in 43.8 minutes (T4), while the R package DAESC-Mix finished the same task in 168.8min (UW Server). For CD8+ T cells, DAESC-GPU-Mix completed the same analysis in 2.3 minutes (A100) and 6.1 minutes (T4), whereas DAESC-Mix used 270.8 minutes (UW Server). For DAESC-GPU, the runtime for CD4+ T cells was longer than that for CD8+ T cells or B cells, which was consistent with expectation due to larger number of cells. However, the runtime of DAESC is shorter for CD4+ T cells than CD8+ and B cells (**Figure 4c**). Upon further investigation, we found that the original DAESC required more iterations to converge for CD8+ T cells and B cells (**Supplementary Figure 4**). DAESC-GPU-Mix identified 15 D-ASE genes between CD4+ naïve vs. central memory T cells (FDR < 0.05). DAESC-GPU-BB identified 6 genes, among which 5 genes overlapped with DAESC-GPU-Mix (**Figure 4d and 4e**). For B cells, DAESC-GPU-Mix and DAESC-GPU-BB identified the same 2 genes (**Figure 4d and Supplementary Figure 5**). Neither DAESC-GPU-BB nor DAESC-GPU-Mix identified significant D-ASE genes between CD8+ naïve vs. central memory T cells. Next, we sought to validate our findings by examining the overlap between the D-ASE genes identified by our methods and the eGenes reported in the original study^17^. For CD4+ T cells, 4 out of 6 genes detected by DAESC-GPU-BB overlapped with the list of eGenes. DAESC-GPU-Mix had the same rate of overlap despite more significant genes, with 10 out of 15 (**Figure 4f**). This finding is consistent with our previous recommendation to use DAESC-GPU-Mix to analyze data with many individuals (e.g., N>20). For B cells, it is difficult to compare the BB and Mix models due to small number of genes (2 genes by both DAESC-GPU-BB and DAESC-GPU-Mix).

**Figure 4.**
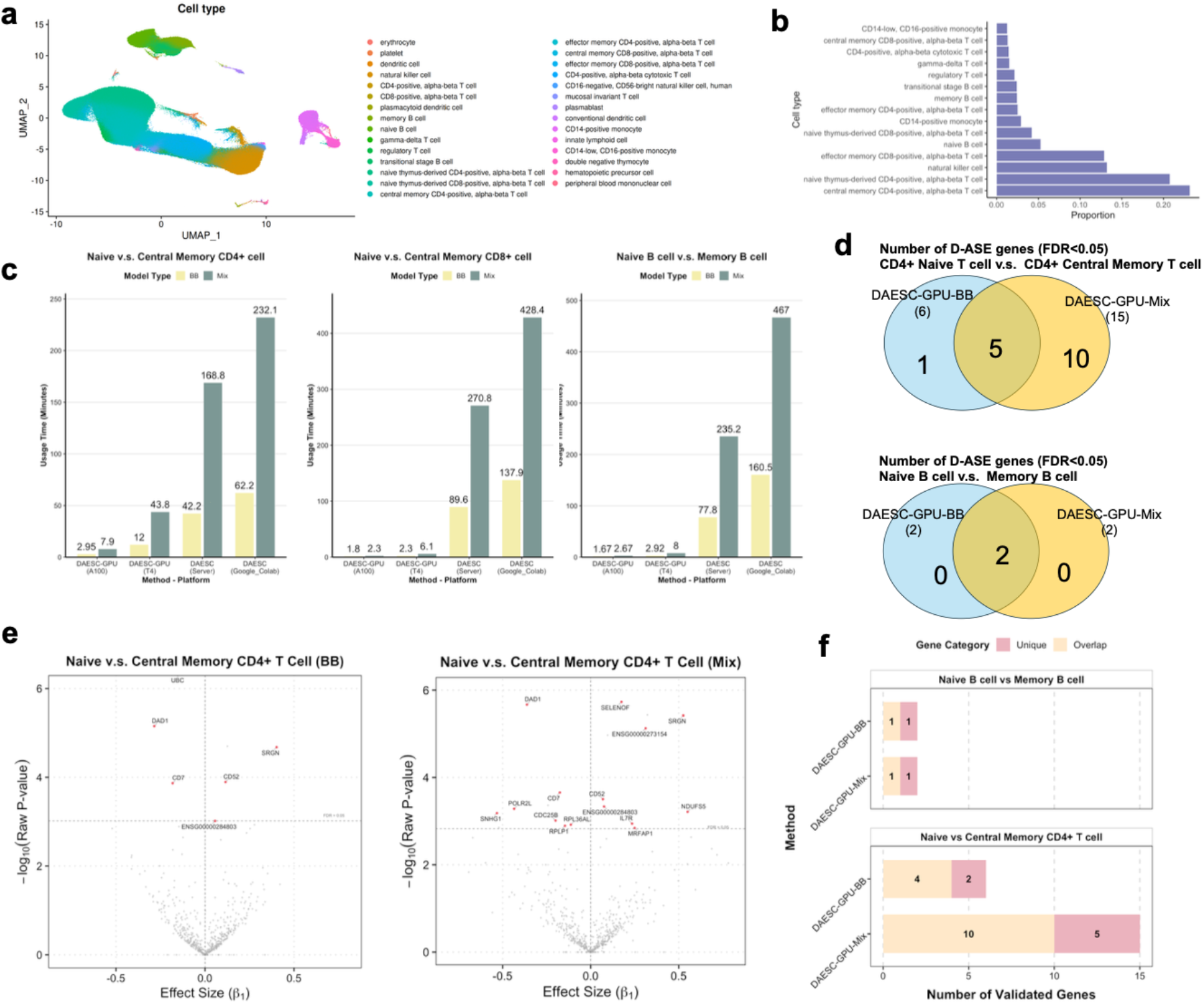
The performance of DAESC-GPU on the OneK1K cohort. (a) UMAP showing all cell types in the OneK1K dataset. (b) Proportion of cell types in the sample (only cell types more abundant than 1% are shown). (c) Runtime comparison between DAESC-GPU (on A100 and T4 GPUs) and the original DAESC R package for differential ASE analyses. (d) Number of significant genes identified by the DAESC-GPU-BB and DAESC-GPU-Mix for CD4+ T cells and B cells (FDR<0.05). No significant genes were found for CD8+ T cells. (e) Volcano plot of differential ASE analysis for CD4+ T cells. (f) Overlap between genes identified by DAESC-GPU and cell-type-specific eGenes reported in the original study as validation evidence. A gene is considered overlapped if it is an eGene in either the naïve or memory cell type.

## Discussion

We presented DAESC+, an integrated software package for single-cell ASE analysis. The preprocessing module, DAESC-P, automates many analysis steps and requires minimal configuration overhead. Compared to SALSA, DAESC-P offers a substantial extension to generate ASE counts from multiplexed scRNA-seq data, making it more widely applicable. The differential analysis module, DAESC-GPU, significantly accelerates the analysis and scales well to large datasets on an entry-level T4 GPU. Its scalability can be extended if a larger number of advanced GPUs are available.

Traditionally, Smart-seq^28,29^ protocols was commonly used for single-cell ASE quantification due to its full-length coverage^22,6,8^. However, it is cost-prohibitive to use Smart-seq to measure many cells and individuals. In contrast, 10x Genomics Chromium has been one of the most popular scRNA-seq technologies due to scalability. However, 10x has a strong bias toward the 3’ or 5’ end of genes and cannot capture the exonic SNPs further inside gene body. This leads to fewer allele-specific reads and higher level of sparsity. Nevertheless, the data we obtained from 10 out of 77 pools of OneK1K were sufficiently powered to detect differential ASE between naïve and memory T cells and B cells. This indicates that 10x can still yield valuable ASE data. While Smart-seq data can be processed with a standard GATK pipeline, 10x data does not have a mature pipeline for ASE extraction. DAESC-P will fill this important gap. In the most complex scenario, DAESC-P can process multiplexed raw reads in SRA format. A full implementation includes downloading SRA files, converting to FASTQ, gather and process cell barcodes, splitting and recombining FASTQ files based on individuals, and downstream steps of SALSA. Users can opt out of some of the steps if their data are not multiplexed or readily in FASTQ format. DAESC-P is organized into several scripts with relatively independent functions, from which users may select as needed.

Although DAESC-P depends on the sequencing technology, DAESC-GPU is a generic software that can be used on single-cell ASE data from any technology, as long as reference and alternative allele read counts are available. We demonstrated that DAESC-GPU improved computational speed many times compared to DAESC for both Smart-seq and 10x data (**Figure 4**). By applying DAESC-GPU to a subset of the OneK1K cohort, we identified genes with cell-type-specific cis-regulatory effects. Those genes were further validated by comparing with previously identified eGenes. These observations indicate that DAESC-GPU can be a candidate tool for analyzing single-cell ASE from other technologies (e.g., long-read sequencing). Long-read sequencing can capture exonic SNPs inside the gene along with isoform information. Comprehensive testing of the software on long-read sequencing is out of the scope of this paper but can be a future direction.

Our study has a few limitations. First, due to the computational demand to process raw sequencing data, we only analyzed 10 pools from the OneK1K dataset. With more computational resources available in the future, we aim to process the entire dataset and conduct a well powered study of differential ASE for immune cells. Second, although DAESC-GPU significantly accelerates the computational speed of DAESC, it does not improve the underlying statistical model. A limitation of this model is its inability to integrate information across multiple discrete cell types. Nevertheless, it remains valuable as a high-performance software package for generic differential ASE analyses. In addition, the runtime advantage of DAESC-GPU in differential analyses between naïve and central memory CD4+ T cells appeared to be less pronounced due to the sparsity of single-cell counts (**Figure 4c**). In such cases, DAESC-GPU organizes the data into large sparse matrices that include all cell-gene combinations with zero counts, while DAESC R package analyzes each gene individually and excludes cells with zero total read counts. This difference narrows their performance gap but DAESC-GPU remains substantially faster.

In conclusion, DAESC+ is a powerful software package for single-cell ASE processing and differential analysis. Application of the software will reveal novel biological insights from the rapidly growing allele-specific datasets.

## Methods

### DAESC-P: Preprocessing module of DAESC+

DAESC-P is an open-source, user-friendly bioinformatics pipeline designed to generate single-cell ASE counts from multiplexed scRNA-seq data. It is comprised of the following steps:

1. The pipeline begins by downloading raw sequencing data (.sra files) from the SRA database using the corresponding accession ID and converts them into FASTQ files using the fastq-dump command from SRA-Toolkit. FASTQ files are renamed based on the naming conventions required by Cell Ranger, including read 1 (R1), read 2 (R2), and index (I1).
2. For each sequencing run in each pool (of individuals), DAESC-P splits multiplexed FASTQ files into individual-specific FASTQ files based on lists of cell barcodes provided by Gene Expression Omnibus (GEO). This step is completed by the BBMap program^30^. Individual-specific single-run fastq files are combined across all runs for each individual.
3. Cell Ranger is used to align each FASTQ file to the reference genome.
4. The remaining steps closely resemble the SALSA pipeline^25^, which includes variant calling using GATK (4.2.0.0) best practices workflow, haplotype phasing using SHAPEIT5^31^, VCF file annotation with GATK Funcotator, alignment bias removal using WASP^21^, and ASE read counting using GATK ASEReadCounter. In contrast to the SALSA pipeline which requires users to run each step separately, DAESC-P packages these steps into a single command and significantly simplifies the process.

Although DAESC-P is built upon SALSA, our pipeline includes two advancements. First, it is tailored to multiplexed scRNA-seq data which pool cells from several individuals in one sample. Direct application of SALSA to such datasets will lead to variant calling errors due to mixing the reads of individuals with different genotypes. If homozygotes are mistakenly treated as heterozygotes, GATK ASEReadCounter can report false monoallelic expression, which dramatically inflate ASE signals. By the splitting and recombining step (step 2), our pipeline accurately calls genotype and counts ASE of each individual. Second, although SALSA is open source, its application requires setting up a complex computing environment. Several steps require downloading large reference datasets, including the GRCh38 reference genome, VCF files documenting known variants in the human genome, GENCODE^32^ and gnomAD^33^. Users must read lengthy documentation of other software (e.g., GATK) to identify and locate the correct reference files. DAESC-P provides all reference data, along with required software packages, into a single file deposited in Zenodo (https://zenodo.org/records/15361243). This significantly reduces the overhead of software configuration, allowing users to conduct data processing with a few simple commands.

### DAESC-GPU: Differential Analysis module of DAESC+

DAESC-GPU is based on our previously developed statistical method for single-cell differential ASE analysis (DAESC)^11^. For a gene or heterozygous transcribed SNP (tSNP), we assume *y*_*ij*_ to be the alternative allele read count of individual *i* and and cell *j*, and *n*_*ij*_ to be the total allele-specific read count. DAESC is based on beta-binomial regression

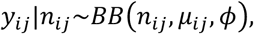

where the average allelic proportion *μ*_*ij*_ is linked to independent variable *x*_*ij*_. DAESC is comprised of a baseline model (DAESC-BB) and a full model (DAESC-Mix). DAESC-BB incorporates individual-specific random effects a_*i*_ in beta-binomial model to account for the sample repeat structure arising from multiple cells measured per individual:

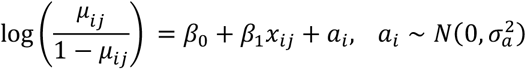

DAESC-Mix further incorporates implicit haplotype phasing using latent variable Z_*i*_ that is either 1 or -1:

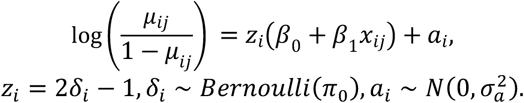

The value of Z_*i*_ depends on whether the alternative allele of the eQTL SNP is on the same haplotype on the alternative or reference allele of the tSNP, which leads to approximately opposite allelic fractions^11^. Variational EM algorithm is used to estimate the unknown parameters and conduct hypothesis testing of differential ASE (H_0_: *β*_1_ = 0).

DAESC-GPU is an accelerated, scalable software for single-cell differential ASE analysis, which overcomes computational limitations of the existing DAESC R package and facilitates the analysis of large-scale data. DAESC-GPU enhances computational efficiency by utilizing Graphics Processing Units (GPUs), which is an electronic circuit that can perform massively parallel computation at high speed. GPUs are integrated in our software through CuPy, an open-source Python library for GPU-accelerated computing using CUDA Toolkit. CuPy provides kernel functions which allow users to write highly specialized and optimized codes running on GPUs.

A number of computational techniques are employed to transform the original R code of DAESC into Python and CuPy. The original R package analyzes one gene at a time. To best leverage the power of GPU computing, our software reframes the algorithm into large matrix operations that can analyze thousands of genes concurrently. An essential technique is a fully vectorized BFGS framework that enables simultaneous optimization of many objective functions (one objective function for each gene). Each step, including gradient updates, inverse Hessian approximations, and search directions, is structured to operate in a parallelized manner across multiple objective functions. Since built-in functions of CuPy are often not available or efficient for our model, DAESC-GPU uses element-wise kernels to integrate multiple mathematical operations into one single specialized function written in C++. For instance, computing the log-likelihood of the model requires the sigmoid function and the difference between digamma functions, for which there are no built-in CuPy functions. By employing custom kernels, these computations are combined into one single function in C++, which maximizes low-level control over GPUs. These features significantly improve the speed of our software. Finally, the efficiency of DAESC-GPU is further enhanced by combining kernel functions and the memory pool feature in CuPy. Computations are performed directly on the original data without requiring additional memory for latent variables.

### Evaluation of DAESC-GPU on endoderm differentiation data

We compared the performance of DAESC-GPU, run on T4 and A100 on Google Colab, and the original DAESC package, run on 8 CPUs from Google Colab (80 GB RAM) and UW Biostatistics Computing Cluster (referred to as “UW Server”, 96 GB RAM), respectively. In silico experiments were conducted on single-cell ASE data from an endoderm differentiation experiment by Cuomo et al^8^. This study profiled gene expression at 4 differentiation times points using Smart-seq2. We obtained SNP-level allele-specific read counts for 105 donors and followed the same preprocessing steps as in our previous paper^11^. In short, we removed SNPs with low mappability and monoallelic expression to reduce potential genotyping error. For each gene, we aggregated SNP-level ASE counts across all SNPs in the exons of the gene and obtained two haplotype-specific counts (hap1 count and hap2 count). We further removed the genes which had non-zero ASE counts in ≤ 20% of the cells. See Qi et al^11^ for more details. After preprocessing, we obtained allele-specific read counts for 4,102 genes and 30,474 cells. We conducted differential ASE analysis along pseudotime to detect dynamically regulated genes during endoderm differentiation. Pseudotime was inferred and provided by the original study^8^.

Next, we evaluated the scalability of DAESC-GPU to larger datasets by simulating additional cells based on the endoderm differentiation data. First, we randomly selected 2,000 genes. To mimic realistic read depth, we used the total read counts (TRC) from real data and kept them fixed in the simulations. To simulate datasets with more cells, we duplicated the real TRC as TRC for the additional cells. For example, we duplicated the 2000 x 30474 TRC matrix once and created a 2000 x 60948 matrix of TRC for 60,948 cells. Larger datasets were simulated by duplicating the TRC matrix more times. Alternative allele counts were simulated using the same procedure as described in the Qi et al^11^, which adopted beta-binomial models incorporating donor-level random effects and haplotype combinations. Model coefficients (*β*_0_, *β*_1_, σ^2^, ϕ) for the simulations were derived by fitting DAESC-BB on the real data and selecting 500 genes with largest |*β*_1_|, as described in Qi et al^11^. The number of cell observations ranged from 30,474 to 1.2 million. To accommodate larger memory requirements, we used an NVIDIA A100 GPU with 40 GB RAM for data analysis. As the number of cells increased, we partitioned the genes into groups to accommodate the limited RAM. For cell counts below 0.35 million, DAESC-GPU 2,000 genes were analyzed simultaneously. For counts between 0.35 to 0.7 million, the genes were divided into two groups and analyzed sequentially. For cell counts between 0.7 and 1.05 million, the genes were split into three groups. For counts exceeding 1.05 million, the genes were divided into four groups. The reported runtime is the sum of runtime of the groups. Due to the rapidly increasing runtime, we restricted the use of DAESC (R package) to datasets with no more than 0.27 million cells. For comparison, we ran the R package on the UW Biostatistics Server with 8 CPUs and 96 GB RAM, and on the Google Colab with 8CPUs and 80 GB RAM.

### OneK1K data analysis

OneK1K is comprised of 982 donors of Northern European ancestry, with scRNA-seq data for ∼1.27 million peripheral blood mononuclear cells (PBMCs). The scRNA-seq data are multiplexed and contain 77 pools, each of which is comprised of 9-18 individuals in multiple runs. We analyzed a subset of the OneK1K cohort (pools 1-10, 104 individuals) to detect differential allelic imbalance between naïve and memory immune cells: naïve versus central memory CD4+ T cells, naïve versus central memory CD8+ T cells, and naïve versus memory B cells. Cell type labels were provided by the original dataset. We used DAESC-P to process the raw sequencing data in SRA format. After completing the pipeline, we obtained a total of 5.7 million nonzero read counts for 28,533 variants across 104 individuals. For each comparison, we first filtered out variants with nonzero read counts in fewer than five individuals. We further filtered out variants with non-zero total expression in less than 1% of all cells. For genes that contained multiple exonic variants, we used the variant with the highest cell-type-specific read depth to represent the gene, ignoring any other variants. Following the preprocessing of the OneK1K dataset, the input data for DAESC-GPU consisted of two large sparse count matrices: one for total read counts and the other for alternative read counts. For benchmarking, we analyzed the data using both the R package and DAESC-GPU. The R package analysis was conducted on the UW Biostatistics servers and Google Colab with 8 CPUs, while DAESC-GPU was run on a single NVIDIA A100 GPU or NVIDIA T4 GPU using Google Colab.

## Availability of Data and Materials

DAESC+ is publicly available on GitHub: https://github.com/ITCUI-XJTLU/DAESC-GPU. Scripts for bioinformatics processing of the OneK1K data are also available on the GitHub site. The endoderm differentiation data by Cuomo et al are available on Zenodo: https://zenodo.org/records/3625024#.YnJ-ivPMKi4. The OneK1K data is available on Gene Expression Omnibus (GEO) via accession number GSE196830.

## Acknowledgements

G.Q. was supported by NIH/NHGRI award K01HG013983.

## Supplementary Figures

**Supplementary Figure 1.**
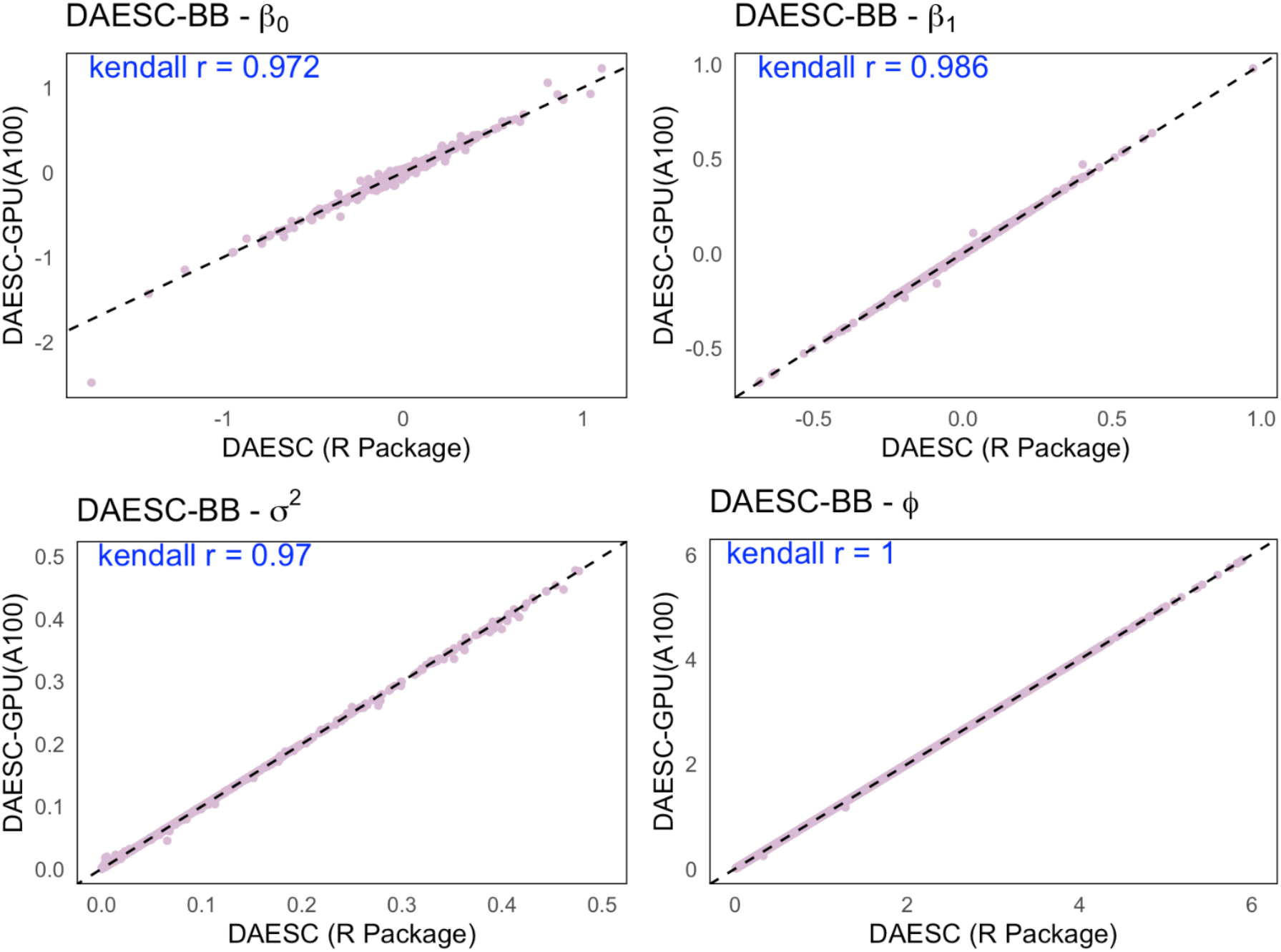
Consistency of parameter estimates between DAESC and DAESC-GPU for the BB model. See Methods for the definition of the parameters. Kendall’s τ is marked in each panel to quantify the consistency. Dashed line is *y* = *x*.

**Supplementary Figure 2.**
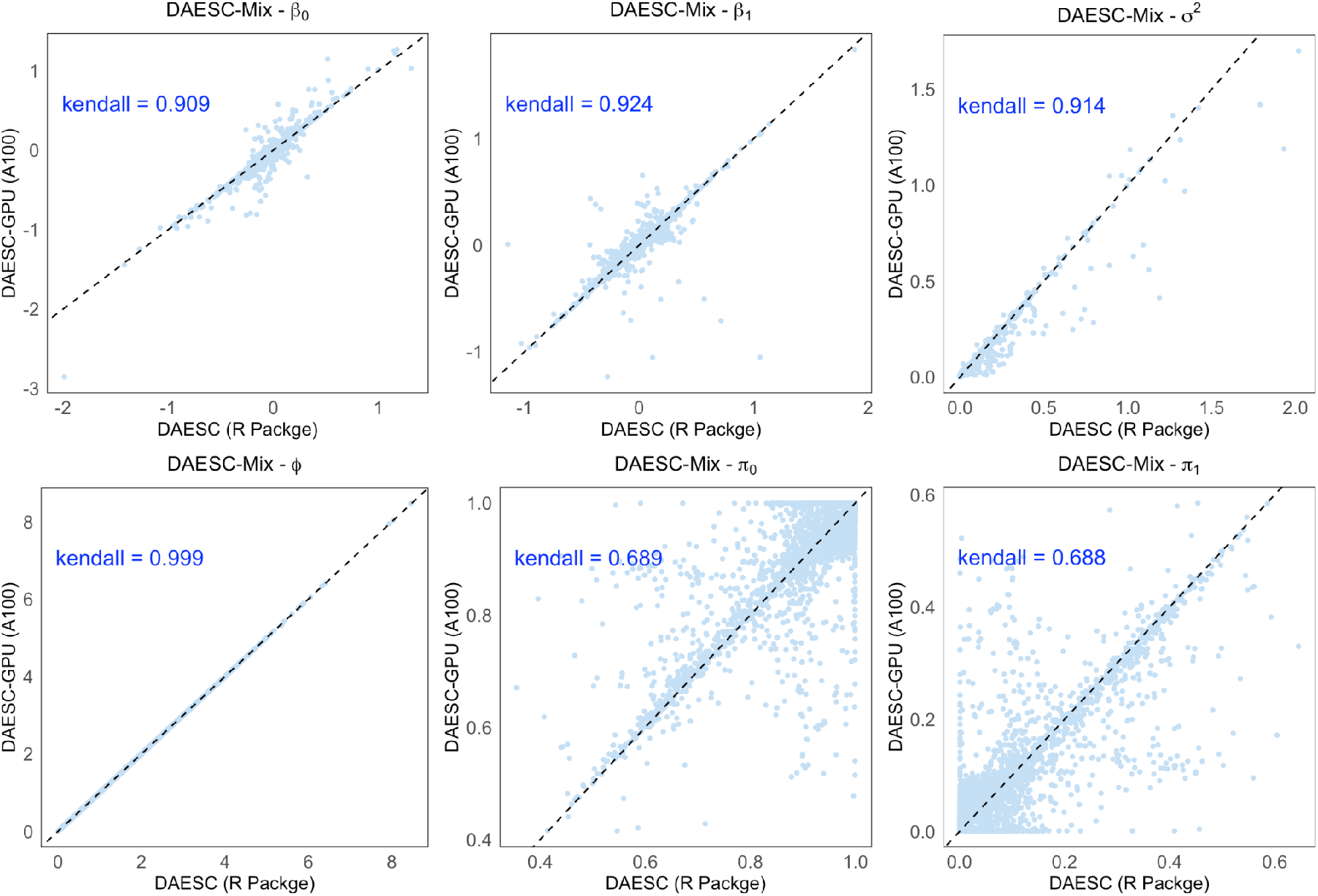
Consistency of parameter estimates between DAESC and DAESC-GPU for the Mix model. See Methods for the definition of the parameters (π_1_ = 1 − π_0_). Kendall’s τ is marked in each panel to quantify the consistency. Dashed line is *y* = *x*.

**Supplementary Figure 3.**
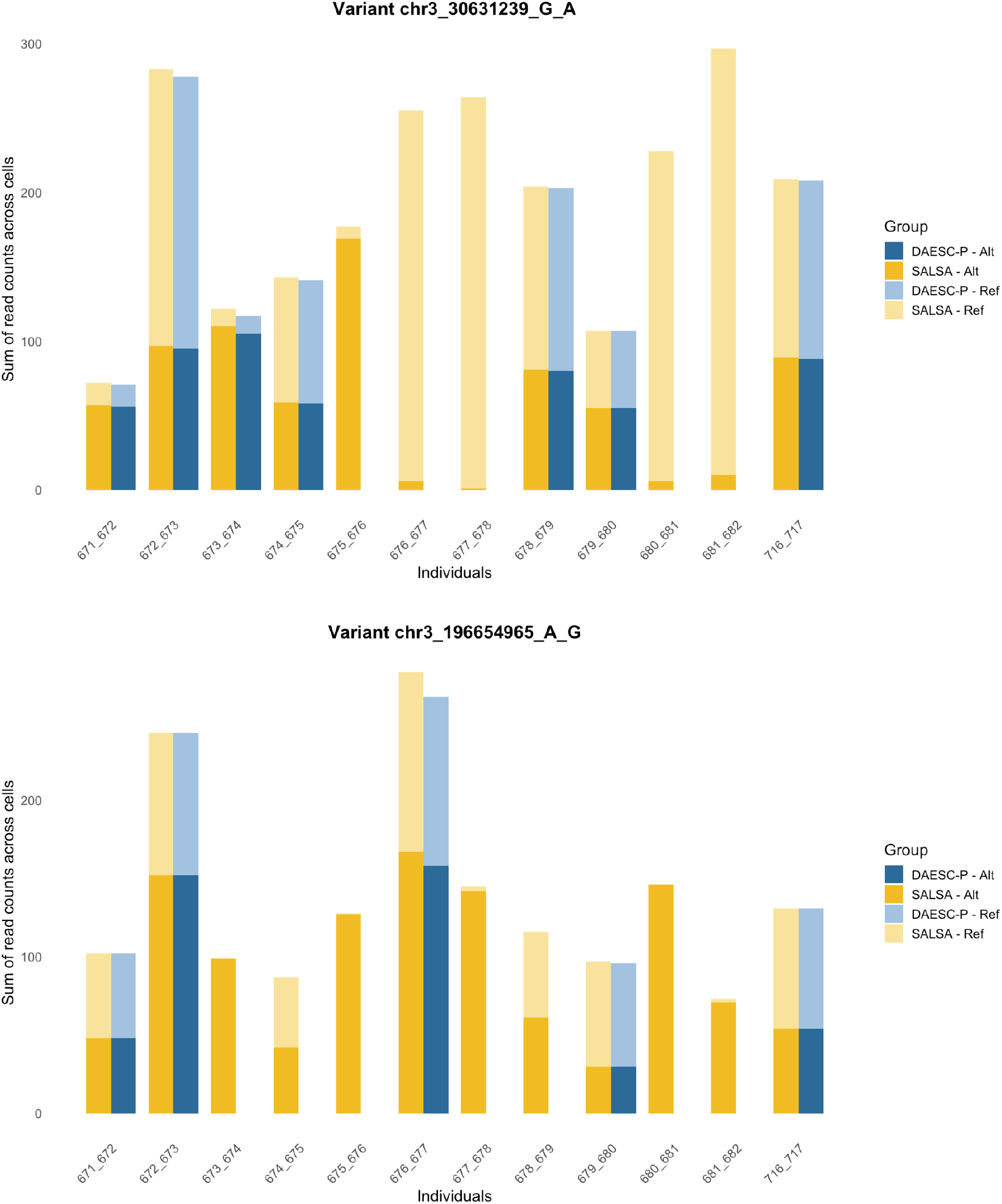
Comparison of ASE counts from SALSA and DAESC-P for two example variants. Reference (ref) and alterantive (alt) allele read counts are summed across all cells for each individual. X-axis indicates the individual ID.

**Supplementary Figure 4.**
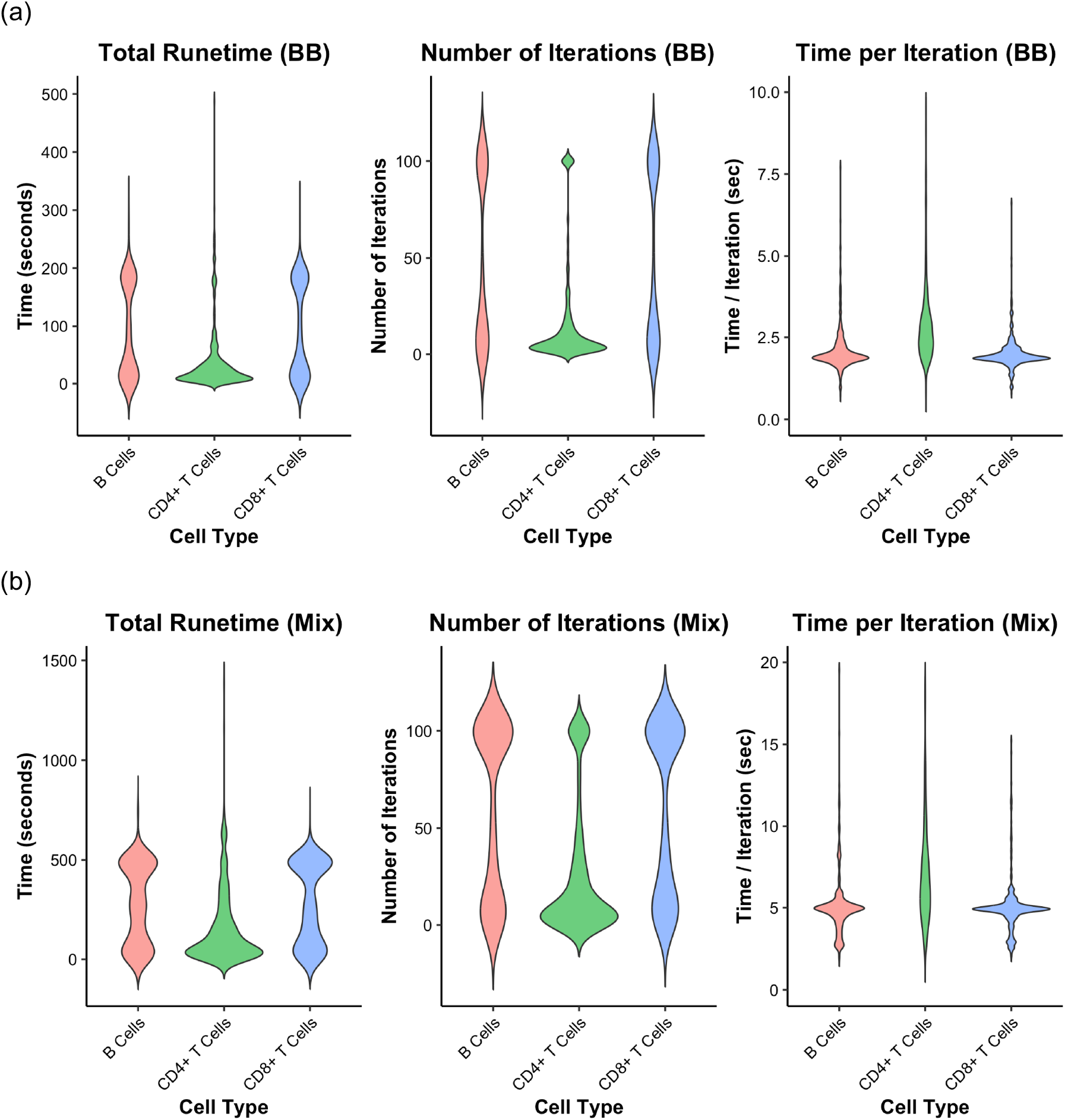
Total runtime, number of iterations and runtime per iteration for DAESC applied to OneK1K. The DAESC R package was run on UW Server. (a) DAESC-BB (b) DAESC-Mix.

**Supplementary Figure 5.**
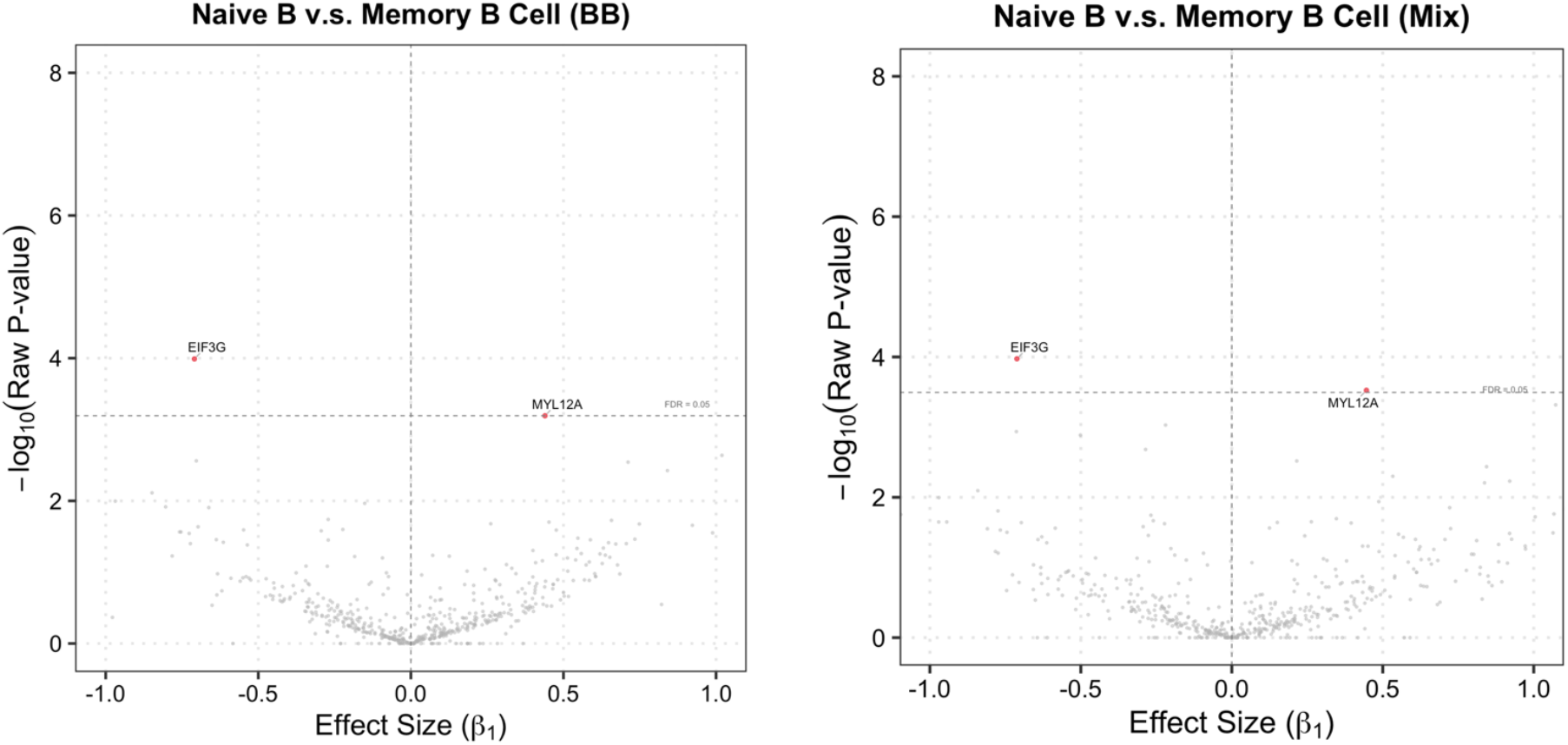
Volcano plot of differential ASE analysis for B cells.

